# Quantitative detection of gut microbial eukaryotes with EukDetect2 reveals global distribution of commensal protists and association with distinct microbial community structure

**DOI:** 10.64898/2026.06.24.734308

**Authors:** Johnathan B. Shih, Chunyu Zhao, Katherine S. Pollard, Abigail L. Lind

**Affiliations:** School of Biological Sciences, Georgia Institute of Technology, Atlanta, GA, USA; Interdisciplinary Graduate Program in Quantitative Biosciences, Georgia Institute of Technology, Atlanta, GA 30332; Gladstone Institute of Data Science and Biotechnology, San Francisco, CA 94158, USA; Department of Epidemiology & Biostatistics, University of California, San Francisco, San Francisco, CA 94158, USA; Chan Zuckerberg Biohub, San Francisco, CA 94158, USA

**Keywords:** metagenomics, fungi, protists, microbial eukaryotes, gut microbiome

## Abstract

Microbial eukaryotes are prevalent members of host-associated and free-living microbial communities, but are routinely excluded from studies of these communities. Existing methods for eukaryote detection from whole metagenome sequencing are limited by contamination of eukaryotic reference genomes and incomplete taxonomic coverage. Our previously published tool EukDetect addressed these challenges using a curated database of universal BUSCO marker genes, but lacked validated quantitative abundance metrics and was built from a limited number of genomes. Here we present EukDetect2, incorporating a database containing 6,948 microbial eukaryotic genomes representing 6,594 unique species, 2,339 of which are newly added since EukDetect version 1, alongside quantitative metrics for estimating absolute and relative abundance of microbial eukaryotes. Using simulated data, we demonstrate accurate abundance estimation, no false positives from bacterial or host-derived reads, and equivalent or greater sensitivity and specificity than alternative taxonomic profiling tools across a range of microbial abundances and community compositions. Applying EukDetect2 across globally distributed human gut microbiome cohorts, we find that *Blastocystis* spp. and *Dientamoeba fragilis* are the most prevalent gut eukaryotes across cohorts, while host-associated fungi are consistently less prevalent than commensal protists. *Blastocystis* abundance is positively associated with a gut microbial community enriched for fiber-fermenting microbes and depleted for pro-inflammatory and industrialization-associated taxa. EukDetect2 provides sensitive, accurate, and quantitative metrics for investigating microbial eukaryotes from metagenomic samples.

## Background

Microbial eukaryotes are ubiquitous members of host-associated and environmental microbiomes, where they function as commensals, mutualists, parasites, and predators of other microbes [1–3]. In the human gastrointestinal tract, protists and fungi engage in complex interactions with the host immune system and with other co-resident microbes. Examples include commensal mouse protists capable of influencing host immune responses and outcompeting gut bacteria for the same nutrient source [4], and commensal fungi such as *Candida* and *Malassezia* species contributing to inflammatory diseases [5,6]. Outside of host-associated microbial communities, fungi function as decomposers with essential roles in global nutrient cycling, and protists act as critical regulators through their functions as phagocytic predators of other microbes and through parasitic associations with invertebrates [1,7–9]. Understanding eukaryotic microbial diversity is therefore critical to the study of microbial communities.

Despite their ecological and clinical relevance, microbial eukaryotes are frequently excluded from microbiome studies. The dominant metabarcoding strategy for characterizing host-associated microbiomes targets the 16S ribosomal RNA gene, which is present only in bacteria and archaea and cannot detect eukaryotes [10]. Eukaryote-specific amplicon approaches targeting the 18S ribosomal gene or the fungal ITS region can identify protists and fungi in dedicated studies, but often miss large groups of taxa, are limited in their taxonomic resolution, and cannot be applied retrospectively to the large body of whole metagenome shotgun sequencing (WMS) data generated by microbiome studies [11–14]. Conversely, WMS captures DNA from all organisms present in a sample, including eukaryotes.

The primary technical barriers to routine eukaryote detection from WMS data are low eukaryotic abundance relative to bacteria and the extensive bacterial contamination present in eukaryotic reference genomes. Eukaryotic genome assemblies frequently contain bacterial-derived sequences, causing spurious alignment to eukaryotic genomes from WMS sequencing, and leading to systemic false positives [15–17]. In addition, host-derived reads can spuriously align to microbial eukaryotic genomes as a result of genome contamination or regions of high sequence similarity, as was recently demonstrated for tumor-associated microbiomes [18].

We previously developed EukDetect to address these challenges [17]. Rather than aligning reads to full eukaryotic genomes, EukDetect aligns reads to a curated database of universal single-copy BUSCO marker genes from which bacterial sequences have been removed, solving the problem of bacterial contamination. However, EukDetect version 1 used a relatively limited number of genomes, relying on the NCBI taxonomy database to choose one representative genome per species. Changes in taxonomy and cryptic species diversity limit the sensitivity of this approach. Additionally, the first version of EukDetect lacked validated absolute or relative abundance metrics.

Here we present EukDetect2, a substantially expanded and updated version of EukDetect that adds new functionality. The EukDetect2 database was constructed from ∼14,600 microbial eukaryotic genomes, which was clustered to include 6,948 representative genomes representing 6,594 NCBI species taxonomy IDs, adding more than 2,000 new species since EukDetect version 1. To construct this database, genomes were clustered at 97% ANI, thus allowing cryptic species-level diversity to persist in the database. EukDetect2 also introduces two quantitative abundance metrics including a proxy for absolute abundance normalized by library size that enables cross-sample and cross-cohort comparisons, and a relative abundance metric expressing the proportional contribution of each detected species to the total eukaryotic fraction of the microbial community. To accommodate the dramatically expanded database and account for more closely related species, which increases off-target alignment [19], we have created a genome and species level disambiguation pipeline that corrects off-target hits and enables accurate quantification. EukDetect2 is both sensitive and specific on simulated sequencing reads from a three-member eukaryotic community, outperforming other eukaryotic classifiers in sensitivity and in detectable taxa.

We characterize the global distribution of human gastrointestinal eukaryotes by applying the EukDetect2 pipeline to nine deeply-sequenced human gut metagenome cohorts spanning countries and lifestyles. We find that the gut protists *Blastocystis* and *Dientamoeba fragilis* are the most common gut eukaryotes globally, with *Blastocystis* reaching near-ubiquity in non-industrialized populations. Conversely, host-associated fungi are less prevalent than gut protists across all groups examined, while dietary fungi are more prevalent than host-associated fungi. Finally, leveraging EukDetect2’s absolute abundance quantification, we find that colonization by the gut protist *Blastocystis* is correlated with a distinct microbial community composition marked by increases in fiber-fermenting bacteria and decreases in mucin-degrading and pro-inflammatory taxa.

## Methods

### Constructing the EukDetect2 database

To construct the EukDetect2 database,14,606 microbial eukaryotic genomes representing 7,944 unique species taxonomy IDs (taxIDs) were downloaded in November 2025. The majority of these genomes were downloaded from GenBank. In addition to the GenBank genomes, we used 76 genomes from the EukProt database [20] representing species without sequenced genomes in the NCBI database, and a re-assembled and filtered *Dientamoeba fragilis* transcriptome (see methods section “*Dientamoeba fragilis* transcriptome re-assembly”). To determine average nucleotide identity (ANI) cluster cutoffs for microbial eukaryotes, all versus all ANI comparisons were performed using skani version 0.3.1 [21]. Clusters of 97% ANI were constructed using galah version 0.4.2 [22]. This resulted in 5,959 clusters of fungal genomes and 1,010 clusters of protist genomes. One representative genome was chosen from each cluster for a total of 6,969 genomes. The *Dientamoeba fragilis* transcriptome was not included in ANI clustering. To determine marker genes in all genomes, marker genes were identified using BUSCO version 6.0.0 using the eukaryota_odb12 database and metaeuk as a gene predictor [23,24]. After running BUSCO, 21 genomes were removed for containing less than 10 BUSCOs. To determine BUSCO genes whose models may result in bacterial sequence contamination, the same BUSCO pipeline was applied to 971 bacterial genomes from the Culturable Genome Reference [25]. We identified 15 BUSCO genes that had calls in these genomes. These BUSCO marker genes were removed from all genomes before the database was compiled.

Genomic regions of BUSCO marker genes were extracted from eukaryotic genomes. These sequences were clustered at a 99% identity threshold using mmseqs2 [26]. Collapsed BUSCO sequences were designed “SSCollapse” if they combined gene duplicates within a single representative genome, “SPCollapse” if they combined genes across multiple species of representative genomes, and “MGCollapse” if they combined genes across multiple genera. To mask repetitive sequence and reduce off-target read alignment, the resulting clustered sequence database was masked with RepeatMasker version 4.1.7 using the FamDB database and the options -qq -noint -norna [27,28].

The final version of the EukDetect2 database after filtering was 6,948 representative genomes, 5,959 of which were fungal (corresponding to 5,663 species taxIDs) and 989 of which were protistan (corresponding to 932 species taxIDs) (Table S1, S2).

### EukDetect2 alignment and post-processing algorithm

To estimate absolute and relative abundances of eukaryotes from metagenomes, sequencing reads are aligned to the EukDetect2 marker database using bowtie2 with parameters optimized for sensitive detection of metagenomes (--end-to-end --very-sensitive, -X 1000) and filtered at a mapping quality also optimized for sensitivity (MAPQ > 10) [19]. To reduce off-target read alignment, reads are only retained if their total aligned length exceeds 80% of the read length, or a minimum of 60 base pairs. Low-complexity reads are filtered by computing the number of unique 4-mers divided by sequence length per read, discarding those below a threshold of 0.5. For paired-end data, a pair is discarded if either mate fails the complexity filter. Duplicate reads are also removed. After alignment and alignment filtering, for each reference marker sequence with one or more mapped reads, EukDetect2 computes read count, coverage, and percent identity using the pysam Python module.

### Taxonomic assignment and disambiguation

Disambiguation is performed both at the genome level, as multiple genomes are assigned to single species taxonomy IDs, and also at the species level. Genome-level disambiguation is performed for species represented by multiple reference genomes in the database. The genome with the greatest number of mapped reads and covered bases is designated the primary genome (or genomes, in the case of ties). Each remaining genome is evaluated as potentially secondary, or representing cross-mapping from the primary, by comparing the overall percent identity between the reads mapping to each genome. If a non-primary genome’s overall percent identity is less than or equal to the primary genome’s, it is classified as a secondary hit. For genomes with fewer than five detected marker genes, a single overlapping BUSCO marker with worse or equal percent identity to the primary is sufficient for being classified as secondary. If the same BUSCO ortholog is present in the primary genome, read counts are then added to the existing primary marker entry. Otherwise, a new marker entry is created for the primary genome, with read count transferring to the primary genome and the read length of the marker in the secondary genome being added to the total marker length of the species. This prevents off-target alignment due to incomplete genomes or incomplete BUSCO annotations.

Species-level disambiguation applies the same logic across all species within a genus. If reads map to multiple species in the same genus, then species with the most reads and covered bases are considered primary. Remaining species are evaluated as possible secondary hits using the same comparisons as for genome-based secondary hits. This step addresses cross-mapping between closely related species. After both rounds of reassignment, a taxon must have reads mapping to at least two distinct marker genes and at least four total reads to be aligned to those genes.

### Relative and absolute abundance estimation of eukaryotes

Absolute abundance is estimated as RPKS (Reads per Kilobase of marker Sequence) and reported as a between-sample comparative metric after normalization for library size, RPKSB (Reads Per Kilobase of marker Sequence per Billion bases sequenced). RPKS is computed as the total reads assigned to a taxon after disambiguation divided by the total length of marker sequences from the primary genome (including any marker sequences that were reassigned during the disambiguation process). Relative abundance (RelEuk) is then computed as each species RPKS divided by the sum of RPKS values detected across all species. RPKS is only reported at the species level.

### *Dientamoeba fragilis* transcriptome re-assembly

A transcriptome of *Dientamoeba fragilis* grown in culture with multiple bacterial species was previously published in 2015 and assembled into 6,595 transcripts [29]. As this is a much lower number of genes than other parabasalid genomes and likely represents an underestimate of the *D. fragilis* transcriptome, we re-assembled the RNA-seq reads and removed likely bacterial sequences. First, RNA-seq reads were downloaded from the SRA under accession SRR2039085 and assembled using Trinity version 2.15.1 [30]. Protein-coding regions were identified using TransDecoder version 5.7.1, resulting in 38,220 protein-coding regions sequences (https://github.com/TransDecoder/TransDecoder). To identify and remove sequences that may have originated from bacteria in the xenic culture with *Dientamoeba fragilis*, we scanned the predicted proteins with the alien index as previously described [31]. Briefly, all proteins (translated CDS regions) were aligned against the clustered NR database using diamond version 2.1.9.163, resulting in 28,586 proteins with a significant hit. The alien index was calculated as *AI* = *bbsO*/*bbsS* − *bbsI*/*bbsS*, where bbsO is the bit score of the best hit to a species outside of the group lineage, bbsI is the bit score of the best hit to a species within the group lineage (skipping self hits), and bbsS is the bit score of the query aligned to itself. The recipient lineage was designated as eukaryotes. Genes with an alien index below 0.1 were considered likely eukaryotic in origin and retained. While this likely excludes genes that are truly present in *D. fragilis* genome, this conservative approach prevents bacterial sequences from entering the EukDetect2 database. Protein-coding regions were clustered at 99% identity to remove duplicates using cd-hit-est version 4.8.1, resulting in 11,404 protein-coding regions [32]. Completeness was assessed with BUSCO version 6.0.0 and the eukaryota_odb12 dataset [23,24]. The protein dataset had 50.4% complete (40.3% single copy, 10.1% multi copy), 10.1% fragmented, and 39.5% missing. This is similar to the BUSCO result for *Trichomonas vaginalis*, the best-annotated parabasalid genome, which is 46.5% complete (41.9% single copy, 4.7% multi copy), 12.4% fragmented, and 41.1% missing, suggesting that the *D. fragilis* proteome presented here represents the same core eukaryotic proteins as other parabasalids.

### Simulated genomic datasets

To determine the susceptibility of the EukDetect2 pipeline to off-target read alignments, we simulated sequencing reads from 971 bacterial genomes from the Culturable Genome Reference Database [25] at 2 million reads per genome, alongside simulating reads from the the human genome (GRCh38), mouse genome (GRCm39), and the model plant *Arabidopsis thaliana* (TAIR 10.1) at 20x genome using art-modern (https://github.com/YU-Zhejian/art_modern). These were chosen to represent common bacterial species in the gut microbiome and common hosts where host-associated microbiota are investigated. Each sample was run through the EukDetect2 pipeline, and for each test case zero microbial eukaryotes were detected. All simulated microbial sequencing and microbial communities in this manuscript were simulated using art-modern (https://github.com/YU-Zhejian/art_modern).

### Public dataset analysis

We analyzed shotgun metagenomic data from nine deeply sequenced publicly available cohorts (Table S3). Seven cohorts represent industrialized populations. Of these, five are from the PREDICT personalized nutrition program (PREDICT1, n = 1,098; PREDICT2, n = 975; PREDICT3 UK, n = 12,340; PREDICT3 US ’21, n = 11,767; PREDICT3 US ’22, n = 8,469), comprising healthy adults recruited in the United Kingdom and United States [33–35]. Additionally, we used a cohort of Israeli adults from a personalized nutrition study (n = 840) [36] and a Dutch cohort enriched for plant-based dietary patterns (n = 150) [37]. To capture gut microbiome compositions from non-industrialized lifestyles, we included a cohort of Hadza hunter-gatherer adults from Tanzania (n = 450) [38] and a cohort from Fiji living in an agrarian lifestyle from (n = 157) [39]. Additional cohort information is available in Table S3.

### *Blastocystis*-bacterial co-abundance analysis

Statistical analyses were performed using R version 4.5.3. For each analyzed public dataset, eukaryotes were removed from the MetaPhlAn4 relative abundance matrix prior to analysis. The remaining bacterial relative abundance matrix was centered log-ratio (CLR) transformed using the R microbiome package (version 1.32.0) to control for the compositional nature of metagenomic relative abundance data. Bacterial taxon abundance was modeled using ordinary least squares linear regression as a function of log1p-transformed Blastocystis RPKSB. Two technical covariates were included in the model: Shannon diversity of the bacterial community and the total sequencing library size. Presence or absence of *Dientamoeba fragilis* was also included as a biological covariate because it is the most frequently co-detected gut protist alongside Blastocystis. Within each cohort, the Benjamini-Hochberg procedure was applied to control the false discovery rate across taxa.

Joint analysis across all cohorts was done with a two-stage approach. In the first stage, replication was assessed across the three largest cohorts (PREDICT3 UK, PREDICT3 US ’21, PREDICT3 US ’22; total n = 32,576). A taxon was considered replicated if it reached Benjamini-Hochberg-adjusted significance (adjusted p < 0.05) in all three cohorts and had a consistent direction of effect. In the second stage, effect sizes were pooled across all seven industrialized cohorts (the three replication cohorts plus PREDICT1, PREDICT2, Zeevi et al, and Shetty et al; total n = 35,634) using random-effects inverse-variance meta-analysis, with between-study variance estimated by restricted maximum likelihood (REML). For each taxon, per-cohort weights were defined as the inverse of within-cohort variance plus estimated between-study variance. The pooled estimate was computed as the weighted mean of per-cohort effect sizes, with the pooled standard error derived from the inverse of the summed weights. Two-sided p-values were computed from the resulting Z-statistics and adjusted for multiple testing using the Benjamini–Hochberg procedure. To prioritize robust cross-cohort ecological associations, pooled results were further restricted to taxa detected in all industrialized cohorts that had concordant effect directions across cohorts (Table S4).

To assess whether *Blastocystis*-associated bacterial taxa show differential prevalence across lifestyle contexts, the prevalence of each taxon in non-industrialized cohorts (Fiji agrarian, Brito_etal and Hadza, Carter_etal) was subtracted from its prevalence in industrialized cohorts (Zeevi_etal, Shetty_etal, PREDICT1, PREDICT2, PREDICT3 UK, PREDICT3 US ’21, PREDICT3 US ’22), yielding a per-taxon prevalence difference. This was computed separately for taxa enriched and depleted in *Blastocystis*-positive samples based on the pooled IVW effect direction. The distribution of these prevalence differences was compared between *Blastocystis*-enriched and *Blastocystis*-depleted taxa using a Wilcoxon rank-sum test. Taxa with positive prevalence differences are therefore more common in non-industrialized populations; taxa with negative differences are more common in industrialized populations.

## Results

### Robust quantification of eukaryotes using an updated database

To construct a curated database of marker genes, we downloaded 14,606 microbial eukaryotic genomes representing 7,944 unique species taxonomy IDs from GenBank in November 2025. An additional 76 genomes not present in GenBank from the eukaryotic genome and transcriptome database EukProt were incorporated. All-versus-all ANI comparison revealed clustering thresholds at 99.5% ANI and at 97% ANI (Figure 1A, Table S1). Genomes clustering at 99.5% identity likely reflect similar strains while genomes clustering at 97% identity likely represent a species boundary. Clustering genomes at a threshold of 97% ANI yielded 5,959 fungal and 989 protist representative genome clusters (Figure 1B, Table S2).

**Figure 1.**
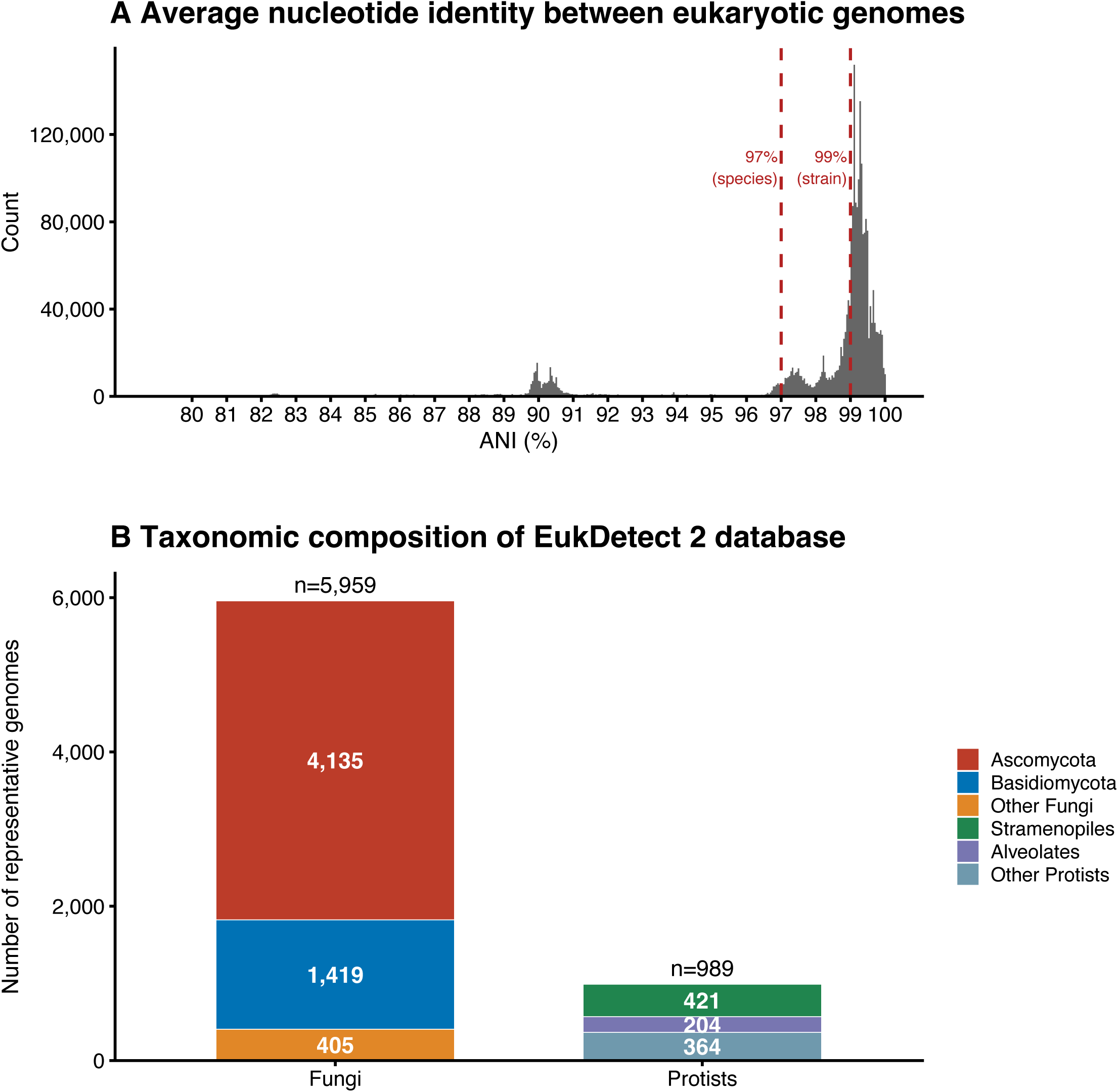
Composition of the EukDetect2 database. (A) Distribution of pairwise average nucleotide identity (ANI) across all-versus-all comparisons of 14,606 microbial eukaryotic genomes downloaded from GenBank. Dashed lines indicate ANI thresholds of 97% (theoretical species boundary) and 99% (theoretical strain boundary). Genomes were clustered at 97% ANI to select representative genomes for the database. (B) Taxonomic composition of the EukDetect2 representative genome database, comprising 5,959 fungal and 989 protist representative genomes.

The EukDetect2 database contains 2,339 new species not in the previous database versions, including 1,919 new fungal and 420 new protistan species (Table S1). In addition, 124 species that were previously represented by only one single genome are now represented by multiple genome clusters, reflecting within-species diversity, cryptic speciation events, or changes in taxonomic classification (Table S2). This includes known species clusters such as the *Aureobasidium melanogenum* and *Aureobasidium pullulans* species complexes, which are represented by 16 and 6 genome clusters, respectively [40].

While most ANI-collapsed genome clusters contained a single species taxonomy ID, 802 clusters had more than one species taxonomy ID (taxID). In some cases, these multi-species clusters correspond to ambiguity in the NCBI taxonomy database (i.e., unnamed species). Each multi-species cluster was manually examined to choose a representative, prioritizing named species, then NCBI-designated representative genomes, then overall assembly quality. Ultimately, 545 species (470 fungi and 75 protists) were represented by a closely related species in the final database rather than by their own genome, representing ∼7% of total species. These comprised closely related species such as members of species complexes in fungi (i.e., the *Trichophyton rubrum* and *Fusarium oxysporum* species complexes), as well as closely related species such as *Leishmania mexicana* and *Leishmania amazonensis*. Species collapsed into a single representative cannot be considered functionally identical. For example, *Aspergillus flavus* and *Aspergillus oryzae* share >99% ANI yet differ in mycotoxin production [41]. However, such species cannot be distinguished from one another in low-coverage metagenomic sequencing data, which is the intended use case of the EukDetect2 pipeline.

### EukDetect2 prevents spurious calls from bacteria and eukaryotic hosts

In many microbial ecosystems, including the human gut microbiome, eukaryotes are lower in abundance than bacteria [13]. Eukaryotic genome assemblies frequently contain bacterial regions either through artifactual contamination during assembly or due to the biological signal of horizontal gene transfer [15]. In either case, these sequences can drive false positive identification of eukaryotes in metagenomes. To determine whether the EukDetect2 database contained bacterial sequences, we simulated 2 million sequencing reads from 971 common human gut bacteria from the Culturable Genome Reference [25] and determined that none of these bacterial genomes produced off-target hits. As with other microbial quantification pipelines designed for host-associated microbiomes, EukDetect2 is designed for sequence libraries where host reads have been removed. However, recent work has demonstrated that fungal detection in metagenomic sequencing of human tissues can arise from human-derived reads [18]. To determine whether the EukDetect2 approach spuriously calls host-derived reads as microbial, we simulated reads from the human genome (GRCh38), mouse genome (GRCm39), and the model plant *Arabidopsis thaliana* (TAIR 10.1) at 20x genome coverage and ran the EukDetect2 pipeline on these samples. For each case, EukDetect2 does not report any eukaryotic microbial taxa as being present, suggesting that eukaryotic host cross-mapping is unlikely to contribute to EukDetect2-reported microbes.

### Accurate metrics to quantify eukaryotes in metagenomics

To enable absolute and relative quantification of eukaryotic members of a microbiome, we developed the metrics RPKSB (Reads Per Kilobase of marker Sequence per Billion bases sequenced), a normalized absolute abundance metric that accounts for the amount of marker gene length present in the EukDetect2 database and is normalized by sequencing library size, and RelEuk, a relative abundance metric that estimates the proportional contribution of each detected eukaryotic species to the total microbial eukaryotic community.

To evaluate the RelEuk metric as an estimate of relative abundance, we simulated sequencing reads from genomes of the human gut protist *Blastocystis* sp. subtype 1, the human-associated fungus *Malassezia restricta*, and the human-associated parasitic gut protist *Giardia duodenalis* at varying genome coverages. In equal coverage simulations at 0.25x and 1x genome coverage, estimated RelEuk values closely matched the expected equal proportions for all three species (Figure 2A). In simulations where one species was present at higher coverage and two species were present at lower coverage, EukDetect2 correctly assigned a higher RelEuk value to the dominant species, with observed values closely tracking expected values across all simulated conditions (Figure 2A).

**Figure 2.**
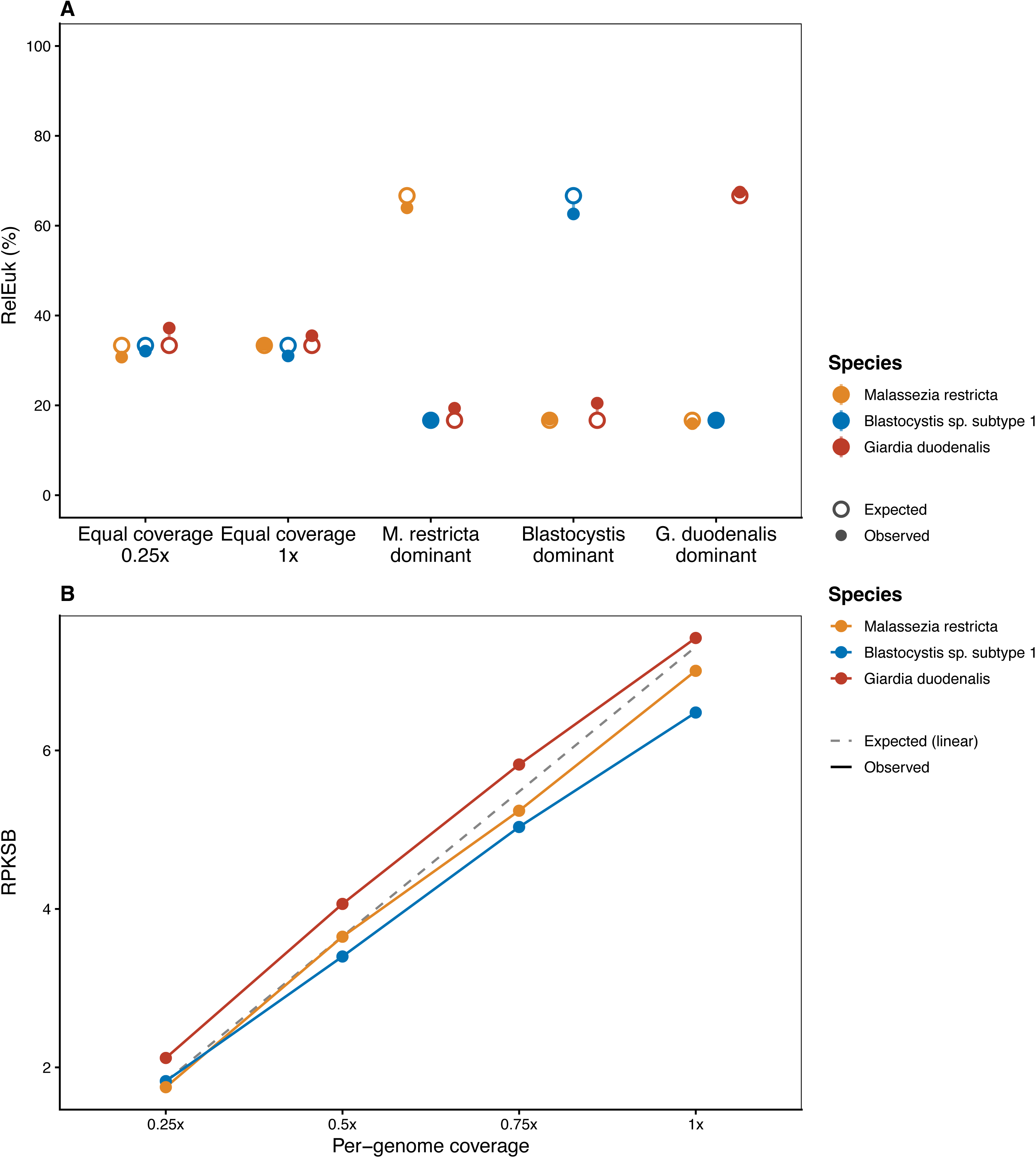
EukDetect2 accurately estimates eukaryotic community composition and abundance in simulated metagenomes. (A) Observed (filled circles) vs. expected (open circles) eukaryotic relative abundance for *Malassezia restricta*, *Blastocystis* sp. subtype 1, and *Giardia duodenalis* across two equal-coverage communities at 0.25x and 1x per genome, and three communities in which one species was simulated at 1x while the others were simulated at 0.25x. (B) RPKSB (Reads per Kilobase of Sequence per Billion bases) scales linearly with per-genome coverage across equal-coverage communities (0.25x–1x). The dashed line indicates expected linear scaling, calculated as the observed median RPKSB at 0.25x scaled proportionally at each successive coverage level.

To evaluate RPKSB as an absolute abundance metric, we compared estimated RPKSB to the per-genome input coverage of each species across equal-coverage simulations. RPKSB scaled linearly for all three species, with observed values closely following the expected linear relationship (Figure 2B). RPKSB is a proxy for overall genome coverage of a species within a particular sequencing library. Attempting to convert this to an estimate of the number of cells of a particular species is complicated by the widely varying DNA content within microbial eukaryotes, driven by differences in ploidy and in nuclei per cell.

### Eukdetect2 is more sensitive and specific than other methods for detecting low-abundance protist and fungi

Several classes of tools are used to detect eukaryotes from shotgun metagenomes, but take different approaches suited to different uses. Methods that align reads to whole eukaryotic genomes or comprehensive nucleotide databases, such as k-mer based classifier Kraken2 and the alignment-based tool CCMetagen, inherit the bacterial sequence contamination widespread in those references [42,43]. Others, such as EukRep and EukFinder, are designed to detect eukaryotic contigs from assemblies; EukRep requires assembled metagenomes and EukFinder classifies reads as eukaryotic in origin for downstream metagenomic assembly prioritization [44,45]. The methods most comparable to EukDetect2 are MetaPhlAn4 and CORRAL, which both classify eukaryotes using curated marker databases [46,47]. CORRAL aligns reads to the EukDetect version 1 database but omits mapping quality filters, instead applying Markov clustering to resolve cross-mapping. The CORRAL taxonomic coverage is therefore identical to EukDetect version 1. MetaPhlAn4 uses lineage-specific markers spanning all domains of life, and its latest database (mpa_vJan25_CHOCOPhlAnSGB_202503, June 2026) contains 489 eukaryotic species (230 fungi, 259 protists) of mixed host-associated and environmental origin.

Across all simulations, EukDetect2 matched or exceeded every tool in sensitivity while producing no off-target eukaryotic calls, and recovered relative abundances closest to expectation (Figure 3, Figure S1). To assess sensitivity and specificity, we simulated sequencing reads across a range of genome coverages for three different eukaryotic species: the gut protists *Blastocystis* sp. subtype 3 and *Entamoba dispar*, and the baker’s yeast *Saccharomyces cerevisiae*. *E. dispar* and *S. cerevisiae* both have close relatives present in all three databases, while *Blastocystis* sp. subtype 3 does not. The lowest coverage simulated was 0.003x for *Blastocystis* and *S. cerevisiae*, and 0.02x for *E. dispar*.

**Figure 3.**
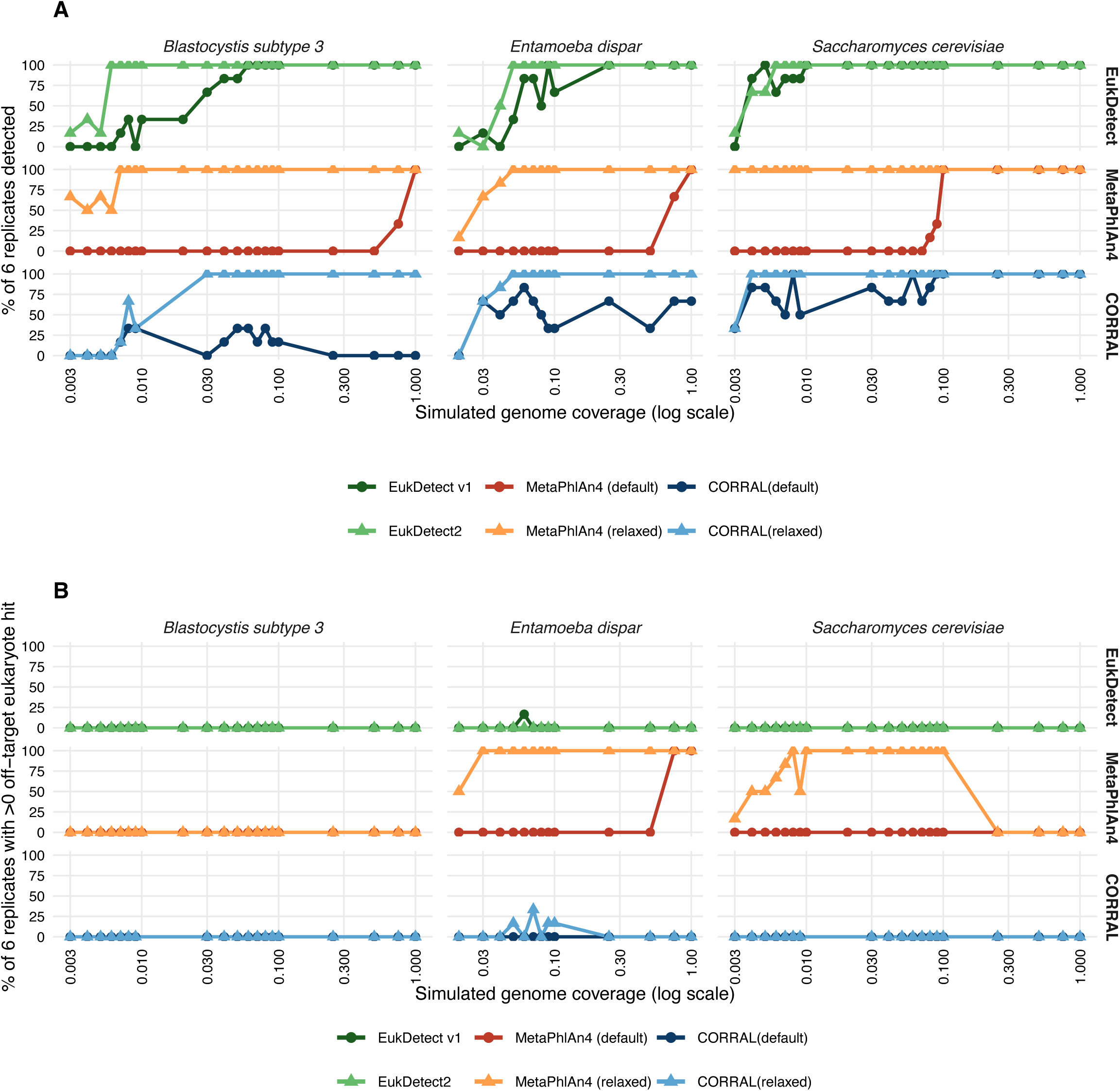
Sensitivity and specificity of eukaryotic profiling tools across simulated genome coverages. Shotgun reads were simulated across a range of coverages for *Blastocystis* subtype 3, *Entamoeba dispar*, and *Saccharomyces cerevisiae* — across six replicates per coverage level. (A) Detection sensitivity, shown as the percentage of replicates in which each target species was detected by each tool. For each species, the x-axis begins at the lowest coverage at which reads were simulated. (B) Off-target detection rate, shown as the percentage of replicates returning one or more non-target eukaryote hits. Results are shown for EukDetect v1, EukDetect2, MetaPhlAn4 (default - stat_q 0.2, and relaxed - stat_q 0 thresholds), and CORRAL (default - PID 97% and relaxed - PID 93% thresholds).

At default settings, neither CORRAL nor MetaPhlAn4 matched the sensitivity of EukDetect2 or version 1 (Figure 3A). Both tools approached EukDetect2’s sensitivity only when their false-positive correcting thresholds were relaxed. Specifically, CORRAL’s percent-identity cutoff was lowered from 97% to 93%, and MetaPhlAn4’s stat_q parameter, which controls which percentiles of marker genes to consider, was set to 0 so that all markers are considered. Both of these adjustments are noted by the authors to increase the false positive rate. We refer to these altered tool settings as CORRAL (relaxed) and MetaPhlAn4 (relaxed).

EukDetect2 was more specific than all other tools tested. No tool reported off-target hits for *Blastocystis* subtype 3, consistent with its lack of close relatives (Figure 3B). For *E. dispar*, however, MetaPhlAn4 (default and relaxed) reported the pathogen *E. histolytica* every time it detected any *Entamoeba*. EukDetect2 did not report any other eukaryote for the *E. dispar* containing samples. CORRAL reported *E. histolytica* in several simulations and EukDetect version 1 did so once. For *S. cerevisiae*, MetaPhlAn4 (relaxed) reported close relatives below 0.03x, the lowest coverage at which MetaPhlAn4 (default) detected the species at all, while CORRAL and both EukDetect versions reported none.

We next evaluated absolute and relative abundance accuracy using simulated mixtures from Figure 2 and the ZymoBIOMICS mock community extracted with the ZymoBIOMICS DNA extraction kit, in which *S. cerevisiae* and *Cryptococcus neoformans* are present at equal DNA proportions [48]. MetaPhlAn4 was assessed only for relative abundance, as it does not estimate absolute abundance. On the simulated mixtures, EukDetect version 1 performed comparably to EukDetect2, with slightly larger deviations from expected values (Figure S1A,B). CORRAL, at its default 97% identity cutoff, estimated abundances poorly across the equal-coverage, *Blastocystis*-dominant, and *Giardia*-dominant communities (Figure S1C); at the relaxed 93% cutoff, it converged on results nearly identical to version 1 (Figure S1D). MetaPhlAn4 consistently underestimated *Blastocystis* sp. subtype 1, producing systematic relative-abundance error (Figure S1E).

On the ZymoBIOMICS sequencing dataset, EukDetect2 detected both species at higher absolute abundance than version 1 (Figure S1F), with a modest gain for *S. cerevisiae* and a substantial one for C. neoformans. Version 1 did not detect *C. neoformans* and yielded 100% relative abundance for *S. cerevisiae*. CORRAL reproduced this version 1 result at default settings but recovered both species when relaxed (Figure S1G). For relative abundance, EukDetect2 came closest to the expected 50:50 ratio, while EukDetect version 1 and CORRAL (default) reported *S. cerevisiae* at 100%, and MetaPhlAn4 and CORRAL (relaxed) deviated modestly (Figure S1H).

This improvement stems from changes in database construction and read filtering. Version 1 represented *C. neoformans* with strain JEC21, since reclassified as *C. deneoformans*; this divergent strain, combined with version 1’s stringent MAPQ > 30 filter, gave poor marker coverage of the ZymoBIOMICS strain. EukDetect2’s ANI clustering circumvents this, increasing sensitivity and robustness to future taxonomic changes.

### EukDetect2 quantifies and identifies a geographic distribution of gut eukaryotes

To evaluate the presence and abundance of common gut eukaryotes, metagenomic sequencing data from nine publicly available cohorts spanning industrialized and non-industrialized populations were processed through the Eukdetect2 pipeline (Figure 4). The commensal protist *Blastocystis* was the most prevalent eukaryote detected across all cohorts. It was near-ubiquitous in non-industrialized populations, reaching 95% prevalence in the Fijian agrarian cohort and 82% in the Hadza hunter-gatherer cohort (Figure 4A). *Blastocystis* was less prevalent in industrialized cohorts, ranging from 50% in the Dutch cohort to as low as 7% in the PREDICT3 US 2022 cohort. Prevalence was consistently lower across US cohorts than others, with the highest US prevalence observed in the PREDICT2 cohort at 12%. *Blastocystis* was also frequently highly abundant when detected, with RPKSB values spanning several orders of magnitude (Figure 4B). Recent work that used a whole-genome alignment based approach to detect *Blastocystis* in all PREDICT cohorts alongside numerous other cohorts found similar, though slightly lower, prevalences of *Blastocystis* in all cohorts [49].

**Figure 4.**
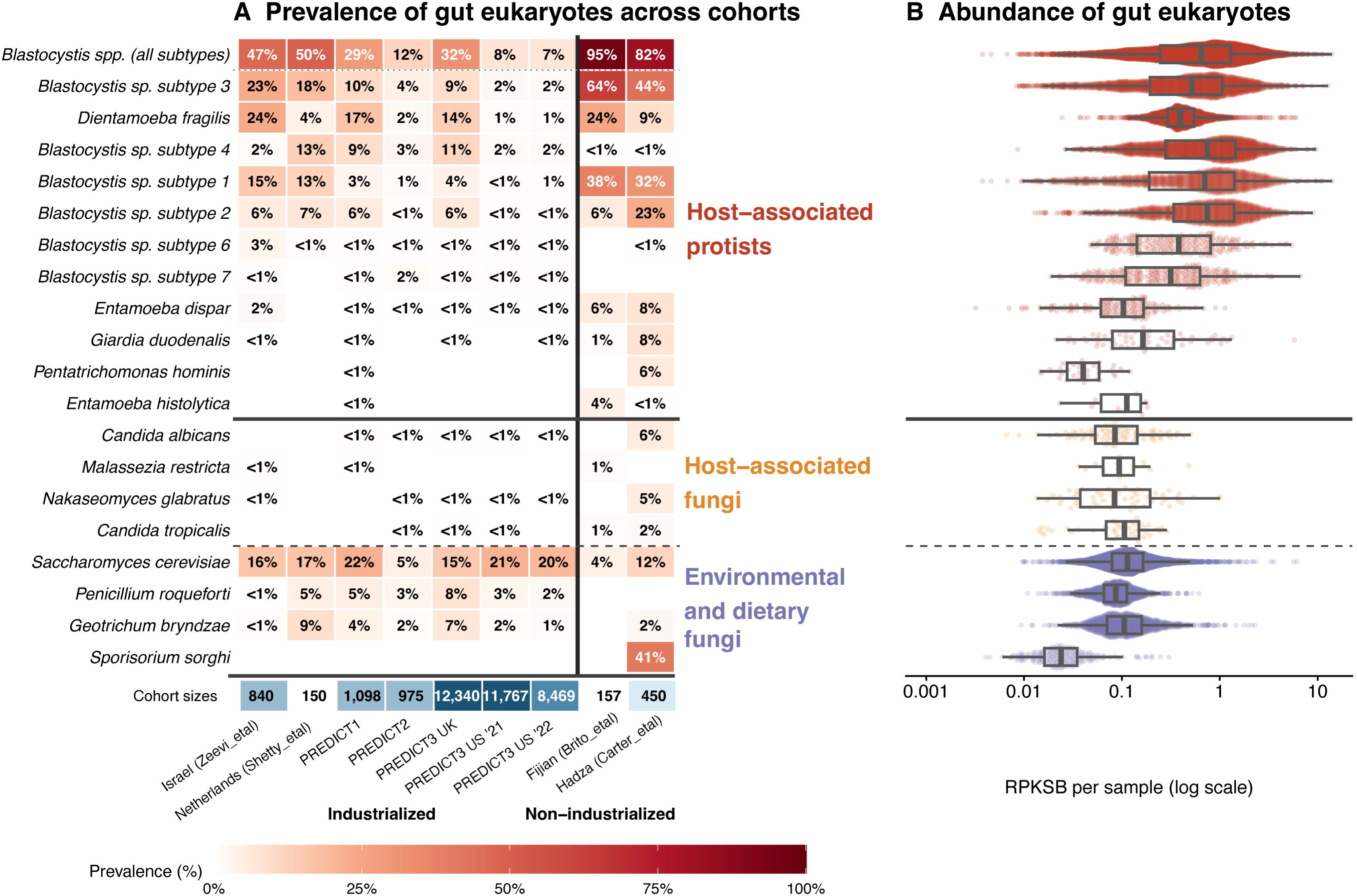
Prevalence and abundance of gut eukaryotes across human cohorts. (A) Prevalence of detected eukaryotic species across nine cohorts spanning industrialized and non-industrialized populations. Cell color and percentage indicate the proportion of samples in which each species was detected. Species are grouped into host-associated protists, host-associated fungi, and environmental and dietary fungi. Solid and dashed lines delineate group boundaries. Cohort sizes are shown below all eukaryotic species. Additional cohort information is in Table S1. In cases where reported cohort sizes in publications and this work differ, this reflects differences in public availability of sequence data. (B) RPKSB distribution for each species across all samples in which it was detected, shown on a log scale. Box plots show median and interquartile range. Points show individual sample values. Zeros are excluded.

The gut protist *Dientamoeba fragilis* was also prevalent, though at lower prevalence than *Blastocystis*, and unlike *Blastocystis* was not substantially increased in prevalence outside of industrialized groups. Prevalence varied between non-industrialized groups; *D. fragilis* was present in 24% of individuals in the Fijian population and 9% of individuals in the Hadza population. Within industrialized groups, the Israeli cohort had the highest frequency at 24% prevalence. *D. fragilis* was least prevalent among US cohorts, at 4% or lower prevalence.

Pathogenic eukaryotes, including *Entamoeba histolytica* and *Giardia duodenalis*, were rare in all cohorts examined and were below 1% prevalence in all industrialized populations. Both species were more frequently observed in non-industrialized settings. *Giardia duodenalis* was present in 8% of Hadza individuals and 1% of Fijian individuals, and *E. histolytica* was present in 4% of Fijian individuals and <1% of Hadza individuals.

Host-associated fungi, including *Candida* and *Malassezia* species, were detected at low prevalence across all cohorts (Figure 4). In contrast, environmental and dietary fungi were more prevalent. *Saccharomyces cerevisiae*, associated with breads and alcoholic beverages, was detected across all cohorts at prevalence ranging from 5% to 22%. Multiple cheese-associated fungi, including *Penicillium roqueforti* and *Geotrichum bryndzae*, were prevalent in European and US industrialized cohorts, consistent with their association with cheese [50]. Fungi are difficult to lyse and can be under-represented in sequencing libraries for this reason. However, the presence of high levels of dietary fungi, including the difficult to lyse filamentous mold *Penicillium roqueforti*, suggests that lysis alone does not explain the lower prevalence of host-associated fungi in metagenomic sequencing datasets, which may instead reflect their true rarity in the gut.

### Eukaryote quantification reveals association with bacterial community structures

Multiple previously published works have established a link between *Blastocystis* carriage and an altered gut microbiome community composition [33,49,51–55]. We assessed the association of other microbes with *Blastocystis* sp., the most prevalent gut eukaryote, using ordinary least squares regression pooling effect sizes across all seven industrialized cohorts, accounting for cohort heterogeneity and adjusting for library size, Shannon diversity of the community, and the presence of the most commonly co-occurring protist, *Dientamoeba fragilis* (see Methods).

In total, 238 microbial taxa were enriched in samples with higher *Blastocystis* loads and 118 microbial taxa were depleted (Table S4). Bacterial and archaeal taxa that were enriched in samples with higher *Blastocystis* load are consistent with a fiber-fermenting and methanogenic gut environment (Figure 5A). *Blastocystis* was positively associated with the presence of the methanogenic archaea *Methanobrevibacter smithii* and *Methanosphaera stadtmanae*, hydrogen-consuming methanogens that remove H₂ generated by primary fermenters [56,57]. Alongside these archaea, enriched taxa included keystone resistant starch and cellulose degraders (*Ruminococcus bromii* and *Ruminococcus champanellensis*) [58,59], primary butyrate producers *(Coprococcus eutactus, Roseburia lenta, Roseburia zhanii,* and *Butyricicoccus intestinisimiae*) [60–62], and Bacteroidaceae capable of complex polysaccharide utilization (*Phocaeicola plebeius*, *Bacteroides pectinophilus*) [63,64]. *Oxalobacter aliiformigenes*, which degrades dietary oxalate from plant foods and is depleted in industrialized populations [65,66], was also enriched.

**Figure 5.**
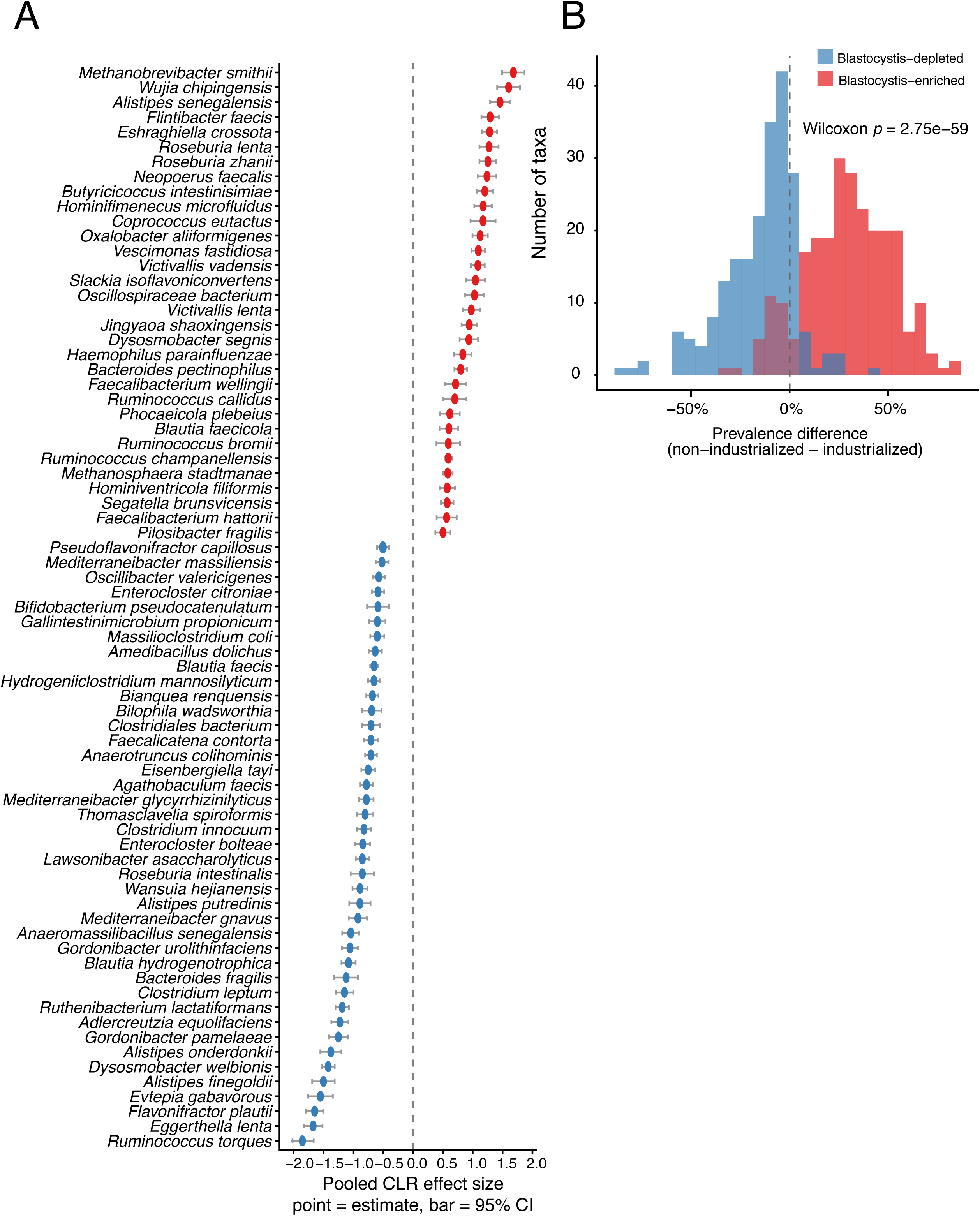
Gut bacterial taxa associated with *Blastocystis* colonization across cohorts. (A) Pooled effect sizes (centered log-ratio, CLR) for bacterial taxa significantly associated with *Blastocystis* presence across cohorts. Red indicates taxa enriched in *Blastocystis*-positive samples; blue indicates depleted taxa. Points show the pooled estimate; bars show 95% confidence intervals. Taxa are ordered by effect size. Only named species taxa are included in this graph, excluding those known by metagenome assemblies alone. A full list of taxa is in Table S4. (B) Distribution of prevalence differences (non-industrialized minus industrialized) for all *Blastocystis*-enriched and *Blastocystis*-depleted taxa. *Blastocystis*-enriched taxa are significantly more prevalent in non-industrialized populations, while depleted taxa are significantly more prevalent in industrialized populations.

Taxa that were depleted in higher *Blastocystis* gut environments include groups that are enriched in non-industrialized groups and are associated with gut inflammation. This includes multiple *Eggerthellaceae* species that transform bile acids and dietary polyphenols, and which in the case of *Eggerthella lenta* have been linked to pro-inflammatory processes in the gut [67,68]. *Mediterraneibacter gnavus* and *Ruminococcus torques*, specialist mucin degraders with links to inflammatory bowel disease [69–71], were negatively associated, as were opportunistic and antibiotic-associated taxa including *Clostridium innocuum* and *Enterocloster bolteae* [72,73].

To assess whether these associations reflect a broader lifestyle signal, we compared the mean prevalence of each taxon in non-industrialized versus industrialized populations across all nine cohorts. Blastocystis-enriched taxa showed significantly higher prevalence in non-industrialized populations relative to industrialized ones, while Blastocystis-depleted taxa showed the opposite pattern (Wilcoxon rank-sum test, p = 2.75 × 10⁻⁵⁹; Figure 5B). Taken together, these findings suggest that Blastocystis colonization is strongly associated with a gut microbial ecology characteristic of non-industrialized, fiber-rich dietary environments, marked by active plant polysaccharide fermentation, methanogenic hydrogen disposal, and the absence of taxa that expand in the context of industrialization.

## Discussion

EukDetect2 represents a significant expansion of detectable eukaryotic species relative to EukDetect version 1, while adding validated absolute (RPKSB) and relative (RelEuk) abundance metrics. By aligning only to curated, decontaminated BUSCO markers and demonstrating in simulations that neither gut bacteria nor host (human, mouse, and *Arabidopsis thaliana*) generate spurious calls, we show that this gain in sensitivity does not come at the cost of specificity. The genome- and species-level disambiguation pipeline makes accurate quantification possible despite the denser, more closely related reference set in the expanded, ANI-clustered database. This clustering also makes EukDetect2 robust to taxonomic reclassification and to the growing number of strains sequenced per species. Benchmarked against its predecessor and against MetaPhlAn4, EukDetect2 captures every species that MetaPhlAn4 detects while additionally recovering low-abundance taxa that MetaPhlAn4 misses, and recovers expected relative abundances in mock communities with less error than MetaPhlAn4 or EukDetect version 1.

Applied across nine globally distributed cohorts, EukDetect2 reveals that the commensal protists *Blastocystis* and *Dientamoeba fragilis* are the most prevalent and abundant gut eukaryotes, far exceeding host-associated fungi, while pathogenic protists like *Entamoeba histolytica* and *Giardia duodenalis* are rare and largely confined to non-industrialized populations. This joins a body of evidence from targeted assays of *Blastocystis* species and *Dientamoeba fragilis* in case-control and general population studies that suggests that these protists constitute normal commensal members of the gut microbiome [74–77]. This does not preclude the possibility of harmful effects in some cases, as many commensal gut bacteria can contribute to disease [78], but argues against treating these protists as primary pathogens.

Host-associated fungi were consistently rare, and only exceeded 1% prevalence in the non-industrialized Hadza population. One possible explanation for the relative rarity of host-associated fungi in gut metagenomic sequencing is that fungi resist common DNA lysis procedures and are therefore under-represented [79]. However, EukDetect2 detected fungi associated with fermented foods, breads, and dairy products more frequently than host-associated fungi like *Candida albicans*. Further, some of the dietary fungi detected are known to be difficult to lyse via standard microbiome DNA extraction protocols. One of the prevalent food-associated fungi detected by EukDetect2 is the blue cheese fungus *Penicillium roqueforti*, which is a filamentous mold. High-yield DNA extraction procedures for filamentous fungi typically include lyophilization of the mycelia followed by grinding in liquid nitrogen [80]. Nevertheless, despite originating from multiple independent studies with different DNA extraction procedures, *Penicillium roqueforti* was detected in European and US cohorts at higher prevalence than any host-associated fungi. Together, these observations indicate that the low signal from host-associated fungi reflects rarity in the gut rather than extraction bias alone.

Our analysis links higher *Blastocystis* abundance to a gut community enriched for fiber-fermenting taxa and depleted for mucin-degrading and industrialization-associated organisms. Additionally, the taxa identified as associated with *Blastocystis* within industrialized cohorts track the broader industrialized–non-industrialized prevalence gradient. We emphasize that this is an association and not evidence of a causal role. *Blastocystis* abundance has been demonstrated to be associated with fiber-rich diets and favorable cardiometabolic markers [49]. Although restricting the meta-analysis to industrialized cohorts and adjusting for colonization by *D. fragilis*, Shannon diversity, and library size mitigates the most obvious confounding factors, other unmeasured dietary or lifestyle factors could drive both *Blastocystis* colonization and the surrounding community structure. Further, differences in sample processing within and between these nine disparate cohorts may also drive taxonomic differences. Whether *Blastocystis* shapes the gut microbiome composition, is a marker for this composition, or both, cannot be resolved from metagenomic surveys. However, this work joins a growing body of evidence that *Blastocystis* presence is associated with an altered gut microbial community [49,51–55]. Work with *Blastocystis* in animal models has been relatively limited due to challenges in establishing long term colonization in mice [81,82]. However, one long-term colonization study in rats demonstrated that colonization by the protist markedly altered the rat gut bacterial community [83]. Further work is needed to determine whether *Blastocystis* colonization alters the gut microbiota directly.

One limitation of the EukDetect2 approach is that it relies on species with available genomes or transcriptomes. Targeted approaches that amplify the 18S region can capture species with only 18S sequences available, though primary efficacy varies and entire eukaryotic groups can be missed due to poor amplification [12,84,85]. With respect to gut eukaryotic diversity, many gut protist species do not have available genome or transcriptome sequences, and therefore our estimates of gut eukaryotic diversity remain a floor, not a ceiling. This impacts detection of eukaryotes in all environments. This includes the amoebozoans *Endolimax nana* and *Entamoeba coli* and the metamonads *Enteromonas hominis, Retortamonas intestinalis,* and *mesnili* [86]. However, the taxonomic coverage of protists continues to improve, and both *Dientamoeba fragilis* and *Pentatrichomonas hominis* have been added to the EukDetect2 database since the first version, alongside increased *Blastocystis* coverage.

## Conclusions

Together, these results establish gut protists as common and abundant members of the human gut microbiome rather than incidental or exclusively pathogenic occupants. By pairing an expanded marker database with validated abundance metrics, EukDetect2 can be applied retrospectively to a large body of shotgun data already generated for microbiome profiling across host-associated and free-living microbial communities.

## Acknowledgements

This work was supported by NHLBI award #HL160862, the Chan Zuckerberg Biohub, and Gladstone Institutes to K.S.P., and by National Institute of Allergy and Infectious Diseases (NIAID) award K22AI173181-02 and Georgia Institute of Technology to A.L.L. Funders had no role in the study design, data collection, analysis, or interpretation, or writing of the manuscript.

## Competing Interests

The authors declare that they have no competing interests.

## Author Contributions

A.L.L. and K.S.P. conceived and designed the study. J.B.S., C.Z., and A.L.L. developed the approach and analyzed the data. J.S., A.L.L., C.Z., and K.S.P. wrote the manuscript. All authors read and approved the final manuscript.

## Availability of data and materials

All sequencing data used in this work was taken from publicly available sources. ZymoBIOMICS mock community data was downloaded from the European Nucleotide Archive under BioProject PRJEB38036. Details for the human gut metagenomic sequencing analyzed in this work, including the public bioproject numbers, is available in Table S1. The assembled *Dientamoeba fragilis* is available on Zenodo at <u>10.5281/zenodo.20818360</u>. EukDetect2 is available on github (https://github.com/allind/EukDetect) and bioconda (https://anaconda.org/channels/bioconda/packages/eukdetect/overview). The EukDetect2 database is available on Zenodo at https://doi.org/10.5281/zenodo.19056625. An archived version of the code used to produce the results in this work is available on Zenodo at <u>10.5281/zenodo.20818571</u>. R code for statistical analysis of bacterial-protist co-abundance is available on github at https://github.com/allind/eukdetect2_analyses.

## Supplementary Tables and Figures

Table S1. Species, genomes, taxonomy IDs, and sources of EukDetect2 database entries.

Table S2. Genome cluster membership of 97% ANI genome clusters in the EukDetect2 database.

Table S3. Public metagenome sequencing cohort information.

Table S4. Table of pooled effect sizes of *Blastocystis* IVW analysis.

**Figure S1**. **Performance of eukaryotic profiling tools on simulated metagenomic communities.** (A–F) Observed (filled circles) vs. expected (open circles) eukaryotic fraction for three simulated species across five community compositions, estimated by EukDetect v2 (A), EukDetect v1 (B), MetaPhlAn4- default (--stat_q 0.2) (C), MetaPhlAn4-relaxed (--stat_q 0) (D), CORRAL-default (PID cutoff 97%) (E), and CORRAL-relaxed (PID cutoff 93%) (F). Line length reflects deviation from expectation. Communities were simulated across two equal-coverage communities at 0.25x and 1x per genome, and three communities in which one species was simulated at 1x while the others were simulated at 0.25x. For EukDetect2, the eukaryotic fraction is the tool’s reported RelEuk metric; for EukDetect v1, RelEuk has been calculated from the existing database; for MetaPhlAn4, it is the species-level relative abundance expressed as a percentage of total eukaryotic relative abundance; for CORRAL, it is the species-level CPM expressed as a percentage of total eukaryotic CPM. (G) RPKSB for EukDetect v2 and v1 on six biological replicates of the ZymoBIOMICS mock community extracted with the ZymoBIOMICS DNA extraction kit [48]. (H) CORRAL-CPM (default and relaxed) on the same community as (G). (I) Eukaryotic fraction across all six tool configurations; dashed line indicates the expected 50% per species. All relative abundances are calculated in the same fashion as A-F.

## Notes

https://github.com/allind/EukDetect

https://doi.org/10.5281/zenodo.19056625

### Competing Interest Statement

The authors have declared no competing interest.

