## Supplementary figures and images for "Quantitative detection of gut microbial eukaryotes with EukDetect2 reveals global distribution of commensal protists and association with distinct microbial community structure"

### Figure S1

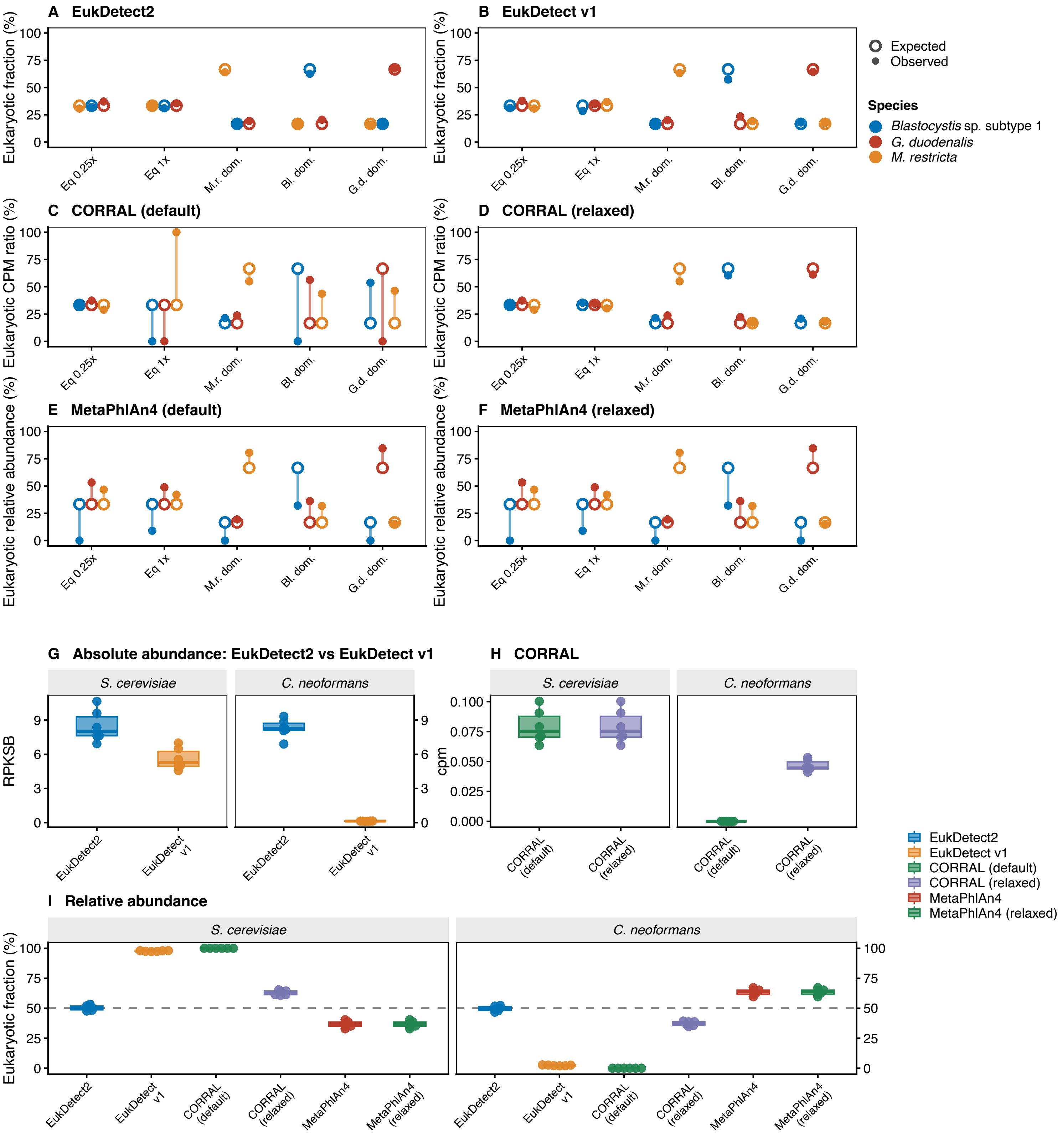
